# GLOBAL LAND-USE AND LAND-COVER DATA: HISTORICAL, CURRENT AND FUTURE SCENARIOS

**DOI:** 10.1101/2021.05.06.442941

**Authors:** Mariana M. Vale, Matheus S. Lima-Ribeiro, Tainá C. Rocha

**Affiliations:** Ecology Department, Federal University of Rio de Janeiro, Brazil; Brazilian Research Network on Global Climate Change (Rede Clima), São José dos Campos, Brazil; National Institute of Science and Technology on Ecology, Evolution and Biodiversity Conservation (INCT EECBio), Federal University of Goiás, Goiânia, Brazil; Biodiversity Department, Federal University of Jataí (UFJ), Goiás, Brazil; Botanical Garden Research Institute of Rio de Janeiro, Rio de Janeiro, Brazil

**Keywords:** Conservation biogeography, ecological niche modelling, macroecology, CMIP6, climate change, deforestation

## Abstract

Land-use land-cover (LULC) data are important predictors of species occurrence and biodiversity threat. Although there are LULC datasets available under current conditions, there is a lack of such data under historical and future climatic conditions. This hinders, for example, projecting niche and distribution models under global change scenarios at different time scenarios. The Land Use Harmonization Project (LUH2) is a global terrestrial dataset at 0.25° spatial resolution that provides LULC data from 850 to 2300 for 12 LULC state classes. The dataset, however, is compressed in a file format (NetCDF) that is incompatible for many analyses and requires layer extractions and transformations that are intractable for most researchers. Here we selected and transformed the LUH2 in a standard GIS format data to make it more user-friendly. We provide LULC for every year from 850 to 2100, and from 2015 on, the LULC dataset is provided under two Shared Socioeconomic Pathways (SSP2-4.5 and SSP5-8.5). We provide two types of files for each year: separate files with continuous values for each of the 12 LULC state classes, and a single categorical file with all state classes combined. To create the categorical layer, we assigned the state with the highest value in a given pixel among the 12 continuous data. LUH2 predicts a pronounced decrease in primary forest, particularly noticeable in the Amazon, the Brazilian Atlantic Forest, the Congo Basin and the boreal forests, an equally pronounced increase in secondary forest and non-forest lands, and in croplands in the Brazilian Atlantic Forest and sub-Saharan Africa. The final dataset provides LULC data for 1251 years that will be of interest for macroecology, ecological niche modeling, global change analysis, and other applications in ecology and conservation.

## Introduction

Land-use and land-cover (LULC) change has been one of the main drivers of environmental change at multiple scales and is currently recognized as an important predictor of anthropogenic impacts and biodiversity threats (Maxwell et al. 2016; Prestele et al. 2016; Gomes et al. 2020; 2021; Rosa et al. 2021). Mapping land-use land-cover (LULC) changes through time is, therefore, important and desirable to predict these threats and propose effective conservation policies (Jetz et al. 2007). LULC is also an important predictor of species’ occurrence and, thus extensively used in ecological and conservation studies (Eyringet al. 2016; Ruiz-Benito et al. 2020; Sobral-Souza et al. 2021). There are several LULC datasets available at a global scale under current conditions, such as the Copernicus (Buchhorn et al. 2020), Global Land Survey, the 30 Meter Global Land Cover, and the GlobeLand30 (Gutman et al. 2013; Pengra et al. 2015; Brovelli et al. 2015), as well as the near historical period, such as the ESA Climate Change Initiative (1992 to 2015), the Finer Resolution Observation, Monitoring of Global Land Cover (1984 to 2011) (Hollmann et al. 2013; Gong et al. 2013) and GCAM (2015-2100) (Chen et al. 2020). These datasets are usually available in standard Geographic Information System (GIS) formats (e.g. TIF or KMZ), routinely used by landscape ecologists, macroecologists, biogeographers and others (Eyringet al. 2016; Ruiz-Benito et al. 2020; Sobral-Souza et al. 2021). However, there is an important gap of historical LULC data covering pre-industrial periods (i.e. older than 1700) and, more importantly, projecting LULC changes into the future. Currently, only two initiatives provide future projections: Global Change Analysis Model (Chen et al. 2020) and Land-Use Harmonization Project (https://luh.umd.edu/data.shtml, Hurtt et al. 2006; 2011; 2020), and only the last one provides a long historical time-series. The absence of compatible dataset across past, present and future scenarios, for example, hinders the use of LULC predictors in projections of ecological niche and species distribution models throughout the time and hamper global change analyses (Escobar et al. 2018).

The recent and robust LULC dataset called Land-Use Harmonization project is part of the Coupled Model Intercomparison Project (CMIP) (https://luh.umd.edu/data.shtml, Hurtt et al. 2006; 2011; 2020), which coordinates modeling experiments worldwide used by the Intergovernmental Panel on Climate Change (IPCC) (Eyring et al. 2016). The data is an input to Earth System Models (ESMs) to estimate the combined effects of human activities on the carbon-climate system. Currently, CMIP datasets are available in NetCDF format, a quite complex file format for most researchers. A few studies used or analyzed the CMIP LULC (Xia & Niu 2020 and references therein), as opposed to CMIP’s climate data already simplified on standard GIS formats available in WorldClim (https://www.worldclim.org/, Fick and Hijmans 2017) and ecoClimate (https://www.ecoclimate.org/, Lima-Ribeiro et al. 2015).

The Land-Use Harmonization project (LUH2) provides the most complete data in term of time-series and scenarios of climate change. The data covers a period from 850 to 2300 at 0.25^°^ spatial resolution (ca. 30 km). The first generation of models (LUH1, Hurtt et al. 2006; 2011) made future land-use land-cover projections under CMIP5’s Representative Concentration Pathways greenhouse gas scenarios (RCPs, see Vuuren et al. 2011), and the current generation of models (LUH2, Hurtt et al. 2020) makes projection under CMIP6’s Shared Socioeconomic Pathways greenhouse gas scenarios (SSP, see Popp et al. 2017). Both provide data on 12 land-use land-cover state classes, including different categories of natural vegetation, agriculture and urban areas. In order to make the global Land-Use Harmonization data more accessible and readily usable, here we filtered, combined and transformed it in standard GIS formats, making the dataset accessible for users with standard GIS skills. Besides providing the Land-Use Harmonization data in regular GIS format at yearly temporal resolution covering 1251 years of past, present and future (from 850 to 2100), we also derived new data based on the existing dataset.

## Methods

We downloaded the 12 land-use land-cover state layers (state.nc) provided in Network Common Data Form (NetCDF) from the Land-Use Harmonization Project (LUH2, https://luh.umd.edu/data.shtml): forested primary land (primf), non-forested primary land (primn), potentially forested secondary land (secdf), potentially non-forested secondary land (secdn), managed pasture (pastr), rangeland (range), urban land (urban), C3 annual crops (c3ann), C3 perennial crops (c3per), C4 annual crops (c4ann), C4 perennial crops (c4per), C3 nitrogen-fixing crops (c3nfx). The “forested” and “non-forested” land-use states are defined on the basis of the aboveground standing stock of natural cover; where “primary” are lands previously undisturbed by human activities, and “secondary” are lands previously disturbed by human activities and currently recovered or in process of recovering of their native aspects (see Hurtt et al. 2006; 2011; 2020 for more details). They were computed using an accounting-based method that tracks the fractional state of the land surface in each grid cell as a function of the land surface at the previous time step through historical data. Because it deals with a large and undetermined system, the approach was to solve the system for every grid cell at each time step, constraining with several inputs including land-use maps, crop type and rotation rates, shifting cultivation rates, agriculture management, wood harvest, forest transitions and potential biomass and biomass recovery rates (see Fig. S1 in the Supplementary Material for details).

To manipulate the NetCDF files, we used the ncdf4 and rgdal packages in R environment (R Core Team 2020, Pierce 2019; Hijmans et al. 2020; Bivand et al. 2021). We also used the Panoply software version 4.8 for quick visualization of the original data (states.nc) (Schmunk, 2017 https://www.giss.nasa.gov/tools/panoply/).

We created two sets of files for each year, the continuous “state-files” and the categorical “LULC-files” (Fig.1, Fig.2 and Fig. S2 of supplemental material). The state-files are the same data provided in the original LUH2 dataset (states.nc), transformed into Tag Image File Format (TIFF) and standardized for ranging from 0 to 1. We built the new LULC-files, also in TIFF format, assigning the highest value among the 12 available states to each pixel. For instance, if the highest value in a given pixel is the forest state value, it was categorically set as a forest pixel. Thus, the LULC-files present categories ranging from 1 to 12, which represents each one of the 12 existing states in the dataset (Table S1 in Supplementary Material). We generated states-files and LULC-files for every year from 850 to 2100 for two greenhouse gas scenarios: an intermediate (SSP2-4.5) and a pessimistic (SSP5-8.5) (see Fig. S2 in Supplementary Material for the workflow to create state files and LULC-files). The SSP2-4.5 scenario, a.k.a “Middle of the Road”, represents a 4.5 W/m^2^ radiative forcing by 2100, where historical development patterns continue throughout the 21^st^ century, susceptibility to societal and environmental changes remains, and greenhouse gas emissions are at intermediate levels. The SSP5-8.5, a.k.a. “Fossil-fueled Development”, on the other hand, represents the upper limit of the SSP scenarios spectrum economic, where social development is coupled with the exploitation of abundant fossil fuel resources, an energy intensive lifestyles, and high levels of greenhouse gas emissions (Popp et al. 2017; Meinshausen et al. 2020; Gatti et al. 2021).

**Figure 1.**
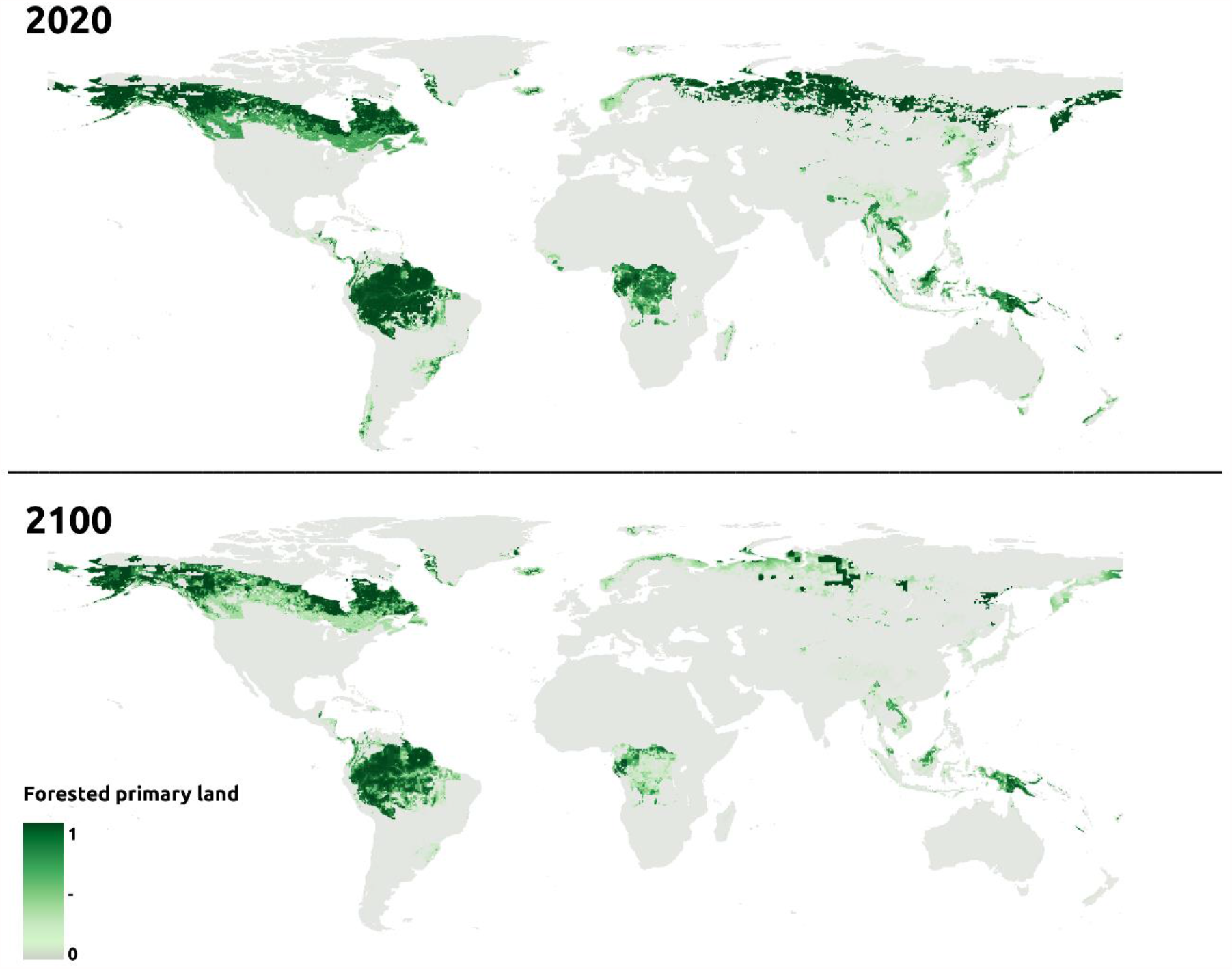
Example of state-files data. Continuous forested primary land state for 2020 (top) and 2100 (bottom) under SSP5-8.5 greenhouse gas scenario, as originally provided by the Land-Use Harmonization (LUH2) project. State values range from 0 to 1, roughly representing the likelihood a pixel is occupied by the land-use land-cover class depicted in the map. All other state-files have the same structure.

**Figure 2.**
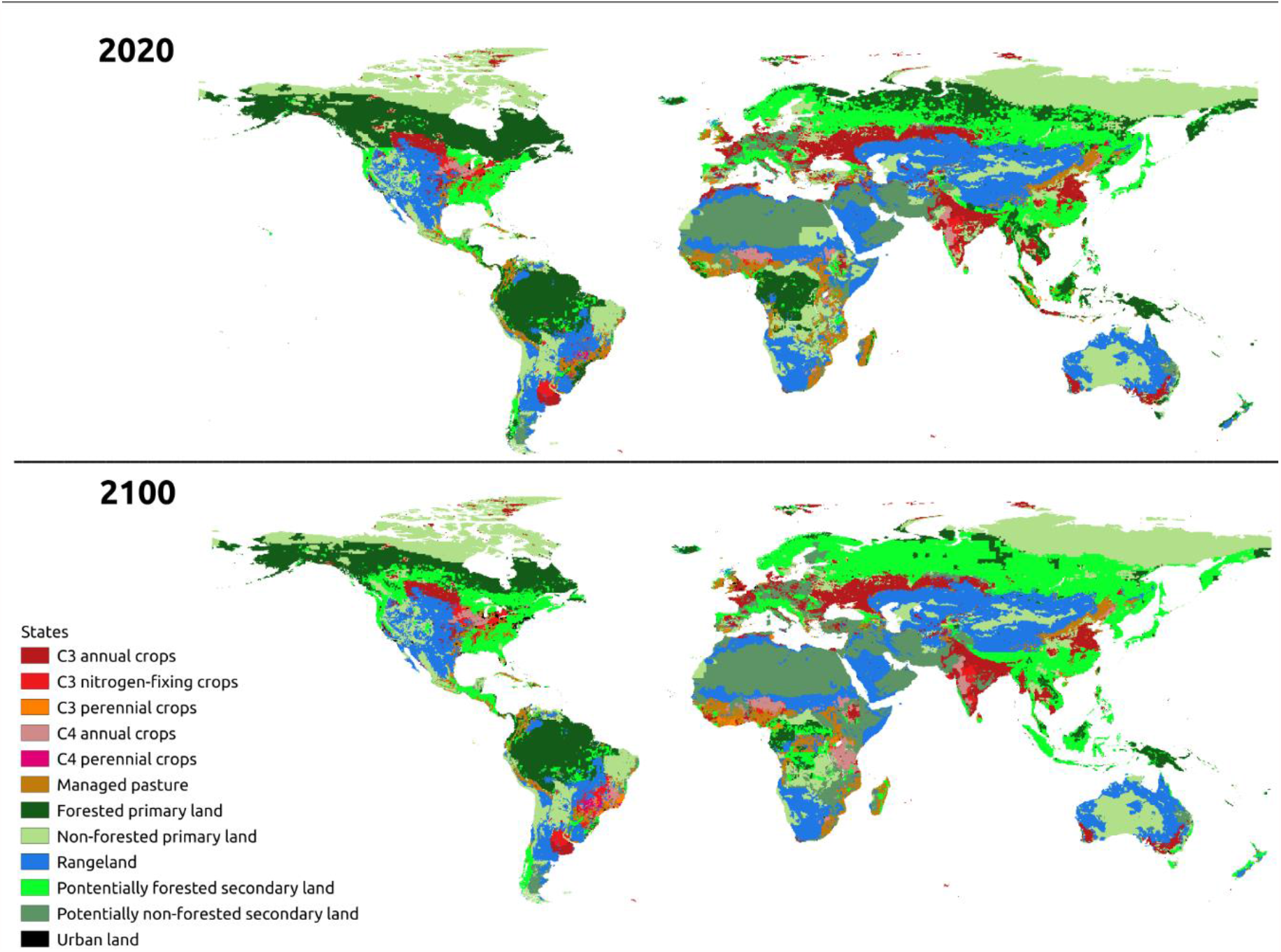
Example of LULC-files data. Categorical LULC for 2020 (top) and 2100 (bottom) under SSP5-8.5 greenhouse gas scenarios, as a result of the combination of the 12 LUH2 original state classes (State-files) into a single map.

We performed an accuracy assessment of our classification for the LULC-files following Olofsson et al.’s (2014) good practices, for the all continents together and for Newton and Dale’s (2001) zoogeographic regions separately. We compared our classified LULC-file for the year 2000 with that of the Global Land Cover SHARE (GLC-SHARE) data, used as the ground truth reference data in the accuracy assessment. The GLC-SHARE was built from a combination of “best available” high resolution national, regional and/or sub-national land cover databases (Latham et al. 2014), and has a finer spatial resolution (1 km) than the LUH2 (30 km). GLC-SHARE has 11 classes that are very similar with those from the LUH2 database: artificial surfaces (01), cropland (02), grassland (03), tree covered areas (04), shrubs covered areas (05), herbaceous vegetation, aquatic or regularly flooded (06), mangroves (07), sparse vegetation (08), bare soil (09), snow and glaciers (10), and water bodies (11). To make the two datasets comparable, we reclassified LUH2 and GLC-SHARE to the following classes: forest, crops, open areas and urban (Fig. 3, Table S1 in Supplementary Material). We also masked-out ice and water areas from GLC-SHARE, as they do not have an equivalent in the LUH2 dataset. Thus, Greenland was removed from analysis and is absent in the LULC-files. We performed the accuracy assessment in QGIS 3.20 through a confusion matrix error, quantifying the commission and omission errors for each class, and then computing three primary metrics: Overall Accuracy (OA), Producer Accuracy (PA) and User Accuracy (UA). We also provide other supplemental metrics, such as Kappa, Allocation Disagreement and Quantity Disagreement using Map Accuracy Tools (Salk et al. 2018) so that users can choose the best metric given their purpose (see supplemental material Accuracies.xlsx).

**Figure 3.**
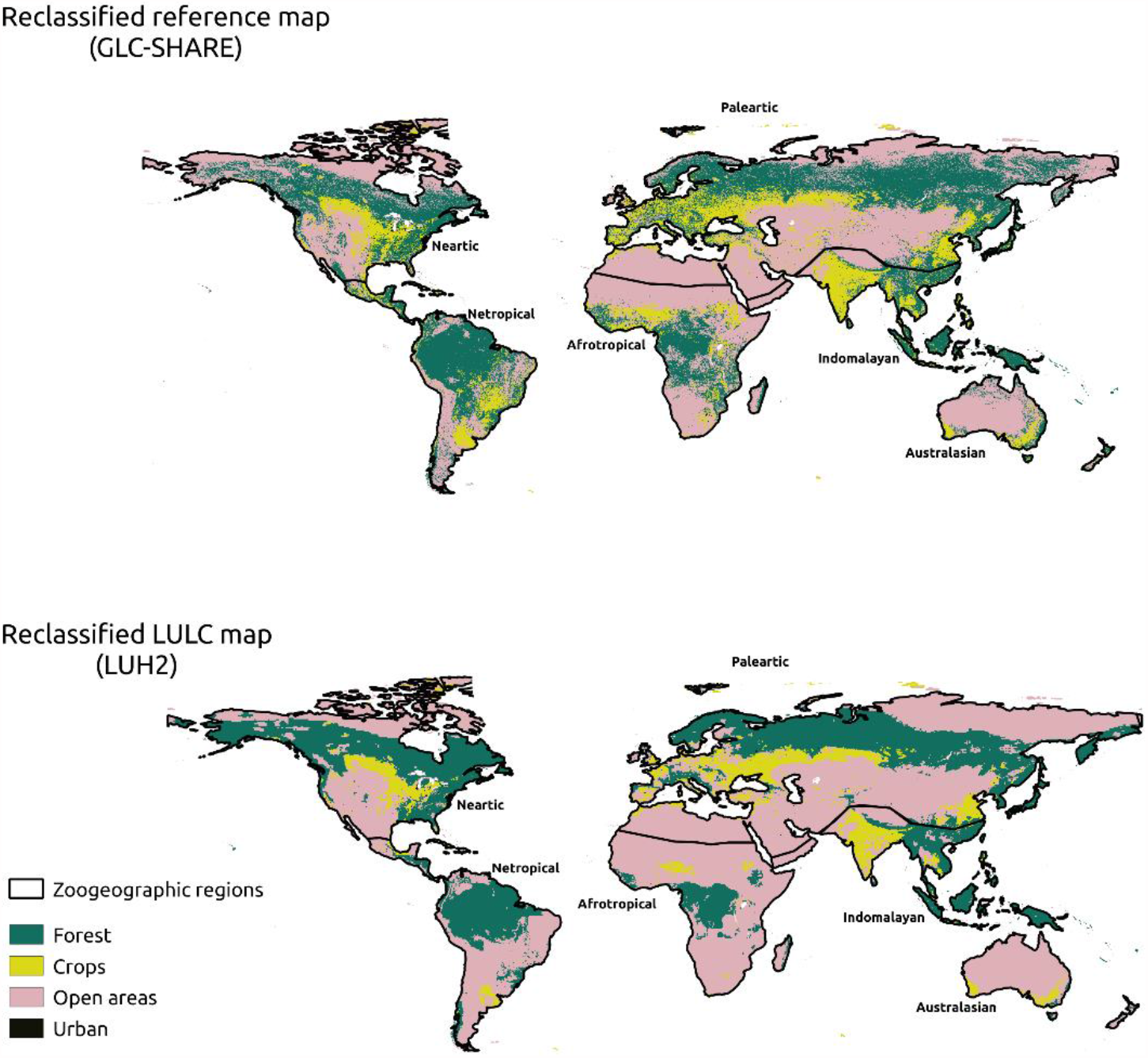
Data used in the accuracy assessment of LULC-files. The accuracy of the classification of the LULC-file (bottom) assessed using the GLC-SHARE as reference data (top). To make the two datasets comparable, both were reclassified to four land-use land-cover states for the year 2000 (see Table 1 for reclassification scheme).

All codes to perform the analysis are available on the GitHub platform (https://github.com/Tai-Rocha/LUH2_Data). The entire resulting dataset is freely available for download at the ecoClimate repository (https://www.ecoclimate.org/), an open database of processed environmental data in a suitable resolution and user-friendly format (Lima-Ribeiro et al. 2015).

## Results

We generated 17.394 files, 16.056 of which are the LUH2 original (continuous data) states files transformed into TIFF (Fig. 1), and the other 1.338 are new (categorical data) LULC-files created by combining the 12 states files (Fig. 2). The LULC-files had good results for most zoogeographic regions and land-use land-cover classes, but not for all (Fig. 3, Table 1). The overall accuracy (OA) was over than 70% for global scale and for most regions, except for the Neotropics, with 65 % overall accuracy. Australasia had the highest OA, with 82% accuracy (see Table 1 and supplemental material S3 for all metrics of accuracy).

**Table 1.**
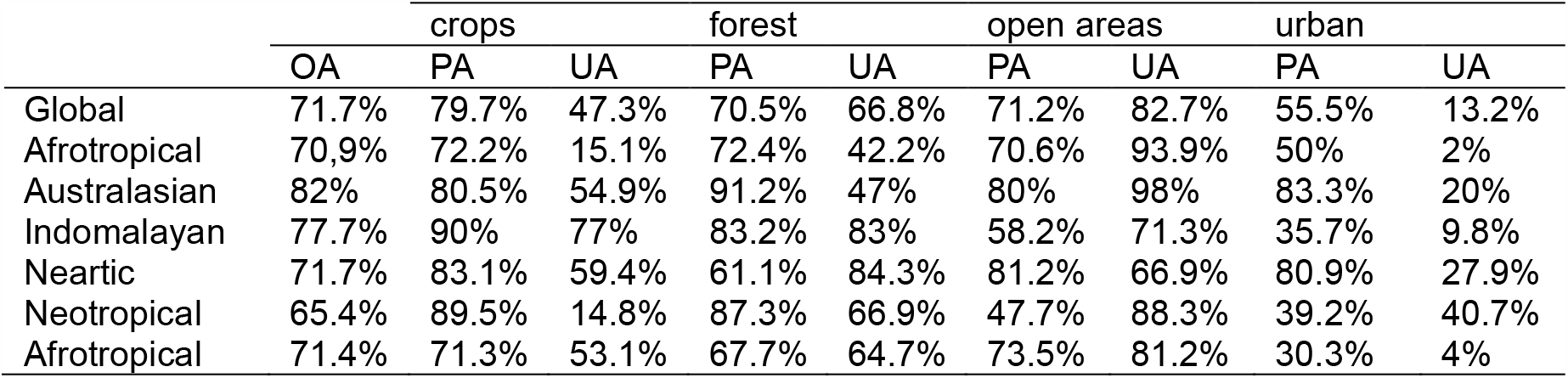
Classification accuracy for LULC classes at global scale and biogeographical regions. OA: overall accuracy; PA: producer accuracy; UA: user accuracy. See the confusion matrix and accuracy metrics in Accuracies.xlsx supplemental file.

The producer accuracy (PA) and user accuracy (UA) for land-use land-cover classes in zoogeographic regions showed some interesting patterns (Table 1 and supplemental tables S3). For crops, there was good PA (71% to 90%) and poor or moderated UA (14% to 59%), except for the Indomalayan region (UA = 77%). Forest had moderate to good PA (61% to 91%) and poor to good UA (42% to 84%). Open area had poor to good PA (47% to 81%), moderate to good UA (71% to 93%). Urban areas had poor to good PA (30% to 83%) and very poor or poor UA (2% to 40%).

The Land-use Harmonized project shows important changes in LULC through time (Fig. 1 and 2), although with no noticeable difference between greenhouse gas scenarios within the same year (Fig. 4). It predicts a pronounced decrease in primary forest, and an equally pronounced increase in secondary forest and non-forest lands (Fig. 4). The decrease in primary forest is particularly noticeable in the Amazon, the Brazilian Atlantic Forest, the Congo Basin and the boreal forests (Fig. 1), coupled with an increase in secondary forest in these regions (Fig. 2). A predicted increase in C4 annual, C3 nitrogen-fixing and C3 perennial crops is especially pronounced in the Brazilian Atlantic Forest and sub-Saharan Africa (Fig. 2). These crops will apparently replace managed pastures in Africa’s Great Lakes region. Finally, there is also a specially pronounced predicted decrease in non-forested primary land (Fig. 4), especially in northern Africa and in the Horn of Africa (Fig. 2).

**Figure 4.**
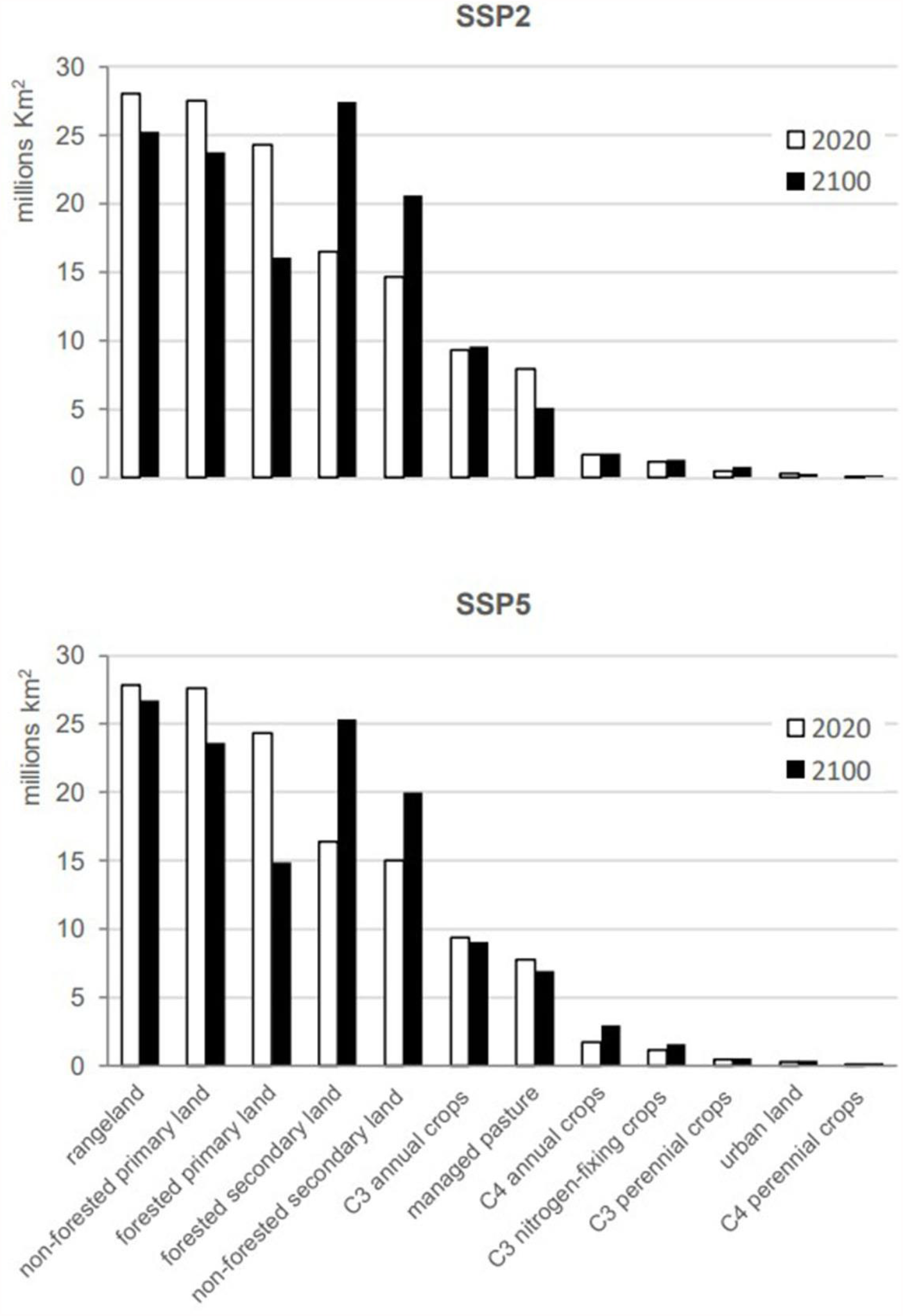
Land-use land cover comparison among years and scenarios. Data for the LULC-files for year 2020 and 2100 for the optimistic (SSP2-4.5, top) and pessimistic (SSP5-8.5, bottom) greenhouse gas scenarios, arranged in decreasing order of class area in 2020.

## Discussion

This data paper is an important contribution in making the Land-Use Harmonization project data more accessible. Here, we provide a global scale LULC dataset with yearly time resolution over a period of 1251 years (from 850 to 2100), and considerable spatial resolution (0.25° long/lat). We contributed not only by transforming the data into standard GIS file format, but also by providing new categorical data on land-use land-cover through a long time period. This LULC database provides support for several research fields in ecology and biodiversity, by disseminating open datasets/open-source tools for a quality, transparent and inclusive science. Our open, ready-to-use and user-friendly database will enable a more robust integration between climate and land-use change within biodiversity science (Titeux et al. 2017; Albert et al. 2020; Hanna et al. 2020).

Given that overall accuracy is still a widely used metric (e.g. Curtis et al. 2018; Gong et al. 2019; Kafy et al. 2021; Liu et al. 2021), our LULC-files provide good quality data (70% to 82% OA), especially for large and coarse scale studies. Besides, we follow the best practices suggested by Olofsson et al. (2014) for validation, considering a reference map with higher quality than the map classification. Validation requires the matching of both maps in terms of classes. Thus, we carefully choose a reference map (GLC-SHARE) that shared similarities with LUH2 in terms of number of classes, which we believe reduced the biases in the reclassification process. In any case, we suggest that users consult Table 1 and supplemental file Accuracies.xlsx for classes’ accuracy at different zoogeographic regions when performing regional analysis.

The most pronounced changes predicted by the Land-use Harmonized project between years 2020 and 2100 are the decrease in primary forest and the increase in secondary forest and non-forested lands (Fig.4, SSP2-4.5 and SSP5-8.5). It is important to note that “primary” refers to intact land, undisturbed by human activities since 850, while “secondary” refers to land undergoing a transition or recovering from previous human activities (Hurrt et al. 2006; 2011; 2020). A major concern regarding the reduction of primary forest is, obviously, habitat loss and associated biodiversity decline, specially of rarer species (Chase et al. 2020; Horta and Santos 2020; Lima et al. 2020), in addition to increased greenhouse gas emissions (Mackey et al. 2020) and likelihood of pandemics associated with viral spillover from wildlife to humans (Dobson et al. 2020). Predicted forest loss is noticeable in the Amazon, Brazilian Atlantic Forest, Congo Basin and boreal forests, especially under the SSP5-8.5 (Fig .1 and Fig. 2), which is in agreement with recent findings.

Svensson et al. (2019) found, for example, a decrease from 75% to 38% in boreal forests between years 1973 and 2013, and Shapiro et al. (2021) showed that over 24 million hectares of forest were degraded in the Congo Basin between years 2000 and 2016. Similar or worse scenarios are happening in the Amazon and Atlantic Forest (Junior et al. 2021; Rosa et al. 2021). This is happening particularly inside Brazil, where recent governmental actions have promoted deforestation and forest fires (Escobar 2019; 2020; Amigo 2020; Silva et al. 2021; França et al. 2021; Qin et al. 2021; Vale et al. 2021), with record deforestation rates in the Amazon (Junior et al. 2021). Although not captured quantitatively at the global analysis (Fig.4), another relevant regional level prediction is the increase in C4 annual, C3 nitrogen-fixing, and C3 perennial crops in the Brazilian Atlantic Forest and sub-Saharan Africa (Fig. 2),. Other studies have similar predictions (Zabel et al. 2019), and the trend is already observed in the Atlantic Forest (Rosa et al. 2021).

The data provided here provides support for several analysis in ecology and biodiversity. The continuous data in the state-files may be particularly useful as predictors in ecological niche modeling (Peterson et al. 2011) or can be combined to species distribution models to reconstruct changes in species distributions (Sofaer et al. 2019; Cazaca et al. 2020). The forested primary land state, for example, can be used to model the distribution of forest-dependent species, as in birds from the Atlantic Forest biodiversity hotspot (Vale et al. 2018). This data has the advantage of being represented in continuous values, as opposed to most discrete land cover data (e.g. all datasets cited in this paper), overcoming the shortcoming of using categorical data as layers in ecological niche modeling (Peterson 2001). More importantly, it allows for the use of land cover data in projections of species distribution under future climate change scenarios. Additionally, the categorical data in the LULC-files can be useful in ecosystem services mapping, especially when working with the widely-used InVEST modeling tool (https://naturalcapitalproject.stanford.edu/software/invest), which is highly dependent on land-use land-cover data (Sharp et al. 2020). The LULC-files can also be used in studies of global change impacts from other perspectives (Mantyka-Pringle et al. 2015; Titeux et al. 2017; Newbold 2018; Clerici et al. 2019; Hong et al. 2019; Jetz et al 2007; Powers and Jetz 2019). Least, but not least, the data can help decision-makers in the construction of evidence based mitigation and conservation policies (Martinez-Fernández et al. 2015; Dong et al. 2018). We hope that the dataset provided here, which is freely available for download at ecoClimate repository (https://www.ecoclimate.org/), can foster the use of land-use land-cover data in many and different fields of study.

## Supporting information

Accuracies

## vii. Acknowledgements

This initiative was possible due to the high-quality data that is maintained and made publicly available by Land-Use Harmonization (https://luh.umd.edu/). We are grateful to Ritvik Sahajpa for helping in the early stages of data acquisition and transformation. This study was developed in the context of the National Institute for Science and Technology in Ecology, Evolution and Conservation of Biodiversity (INCT EECBio, CNPq Grant n 465610|2014-5, FAPEG 201810267000023) and the Brazilian Network on Global Climate Change Research (Rede CLIMA). MMV and ML-R received support from the National Council for Scientific and Technological Development (CNPq Grants no. 304309/2018-4 and 301514/2019-4, respectively) and TR from CAPES (Coordination for the Improvement of Higher Education Personnel - Grant No. 88887.373031/2019-00) and Open Life Science (OLS-2).

