## Supplemental material for "GLOBAL LAND-USE AND LAND-COVER DATA: HISTORICAL, CURRENT AND FUTURE SCENARIOS"

**Supplementary Material**

Table S1. Reclassification of the reference and target dataset for accuracy assessment analysis. The reference (GLC-SHARE) and classified (LUH2) data were reclassified into four common classes: (1) forest, (2) crops, (3) open areas, and (4) urban. Classes absent in the focus dataset were masked-out.

| **original class code** | | | **original**  **class name** | **code (name) after reclassification** |
| --- | --- | --- | --- | --- |
| GLC-SHARE | | | | |
| 1 | | Artificial surfaces | | 4 (urban) |
| 2 | | Cropland | | 2 (crops) |
| 3 | | Grassland | | 3 (open areas) |
| 4 | | Tree covered areas | | 1 (forest) |
| 5 | | Shrubs covered areas | | 3 (open areas) |
| 6 | | Herbaceous vegetation, aquatic or regularly flooded | | 3 (open areas) |
| 7 | | Mangroves | | 1 (forest) |
| 8 | | Sparse vegetation | | 3 (open areas) |
| 9 | | Bare soil | | 3 (open areas) |
| 10 | | Snow and glaciers | | masked-out |
| 11 | | Water bodies | | masked-out |
| LUH2 | | | | |
| 1 | C3 Annual Crops (c3ann) | | | 2 (crops) |
| 2 | C3 Nitrogen-Fixing Crops (c3nfx) | | | 2 (crops) |
| 3 | C3 Perennial Crops (c3per) | | | 2 (crops) |
| 4 | C4 Annual Crops (c4ann) | | | 2 (crops) |
| 5 | C4 Perennial Crops (c4per) | | | 2 (crops) |
| 6 | Managed Pasture (pastr) | | | 3 (open areas) |
| 7 | Forested Primary Land (primnf) | | | 1 (forest) |
| 8 | Non-Forested Primary Land (primn) | | | 3 (open areas) |
| 9 | Rangeland (range) | | | 3 (open areas) |
| 10 | Potentially Forested Secondary Land (secdf) | | | 1 (forest) |
| 11 | Potentially Non-Forested Secondary Land (secdn) | | | 3 (open areas) |
| 12 | Urban Land (urban) | | | 4 (urban) |

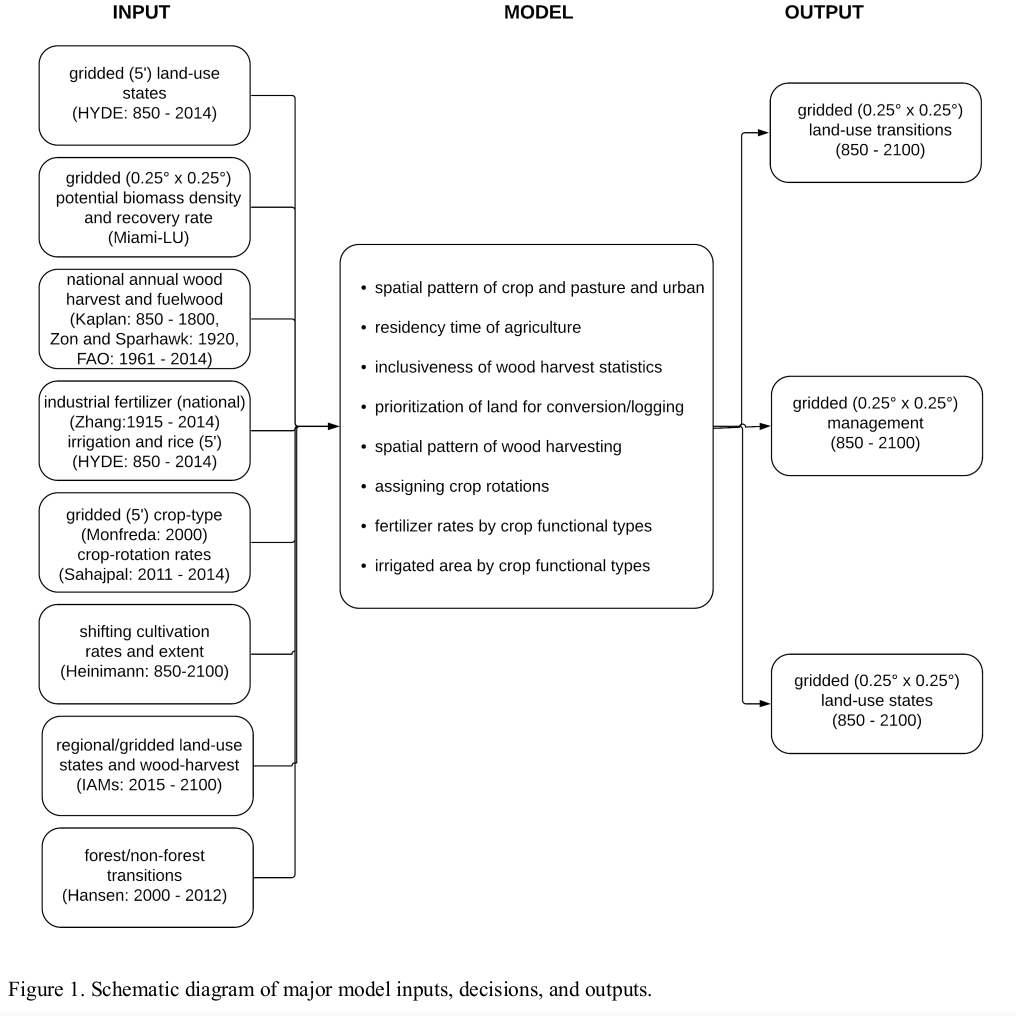

Figure S1. Framework to generate land use and land cover states of Land-Use Harmonization (LUH2, Hurtt et al . 2020)

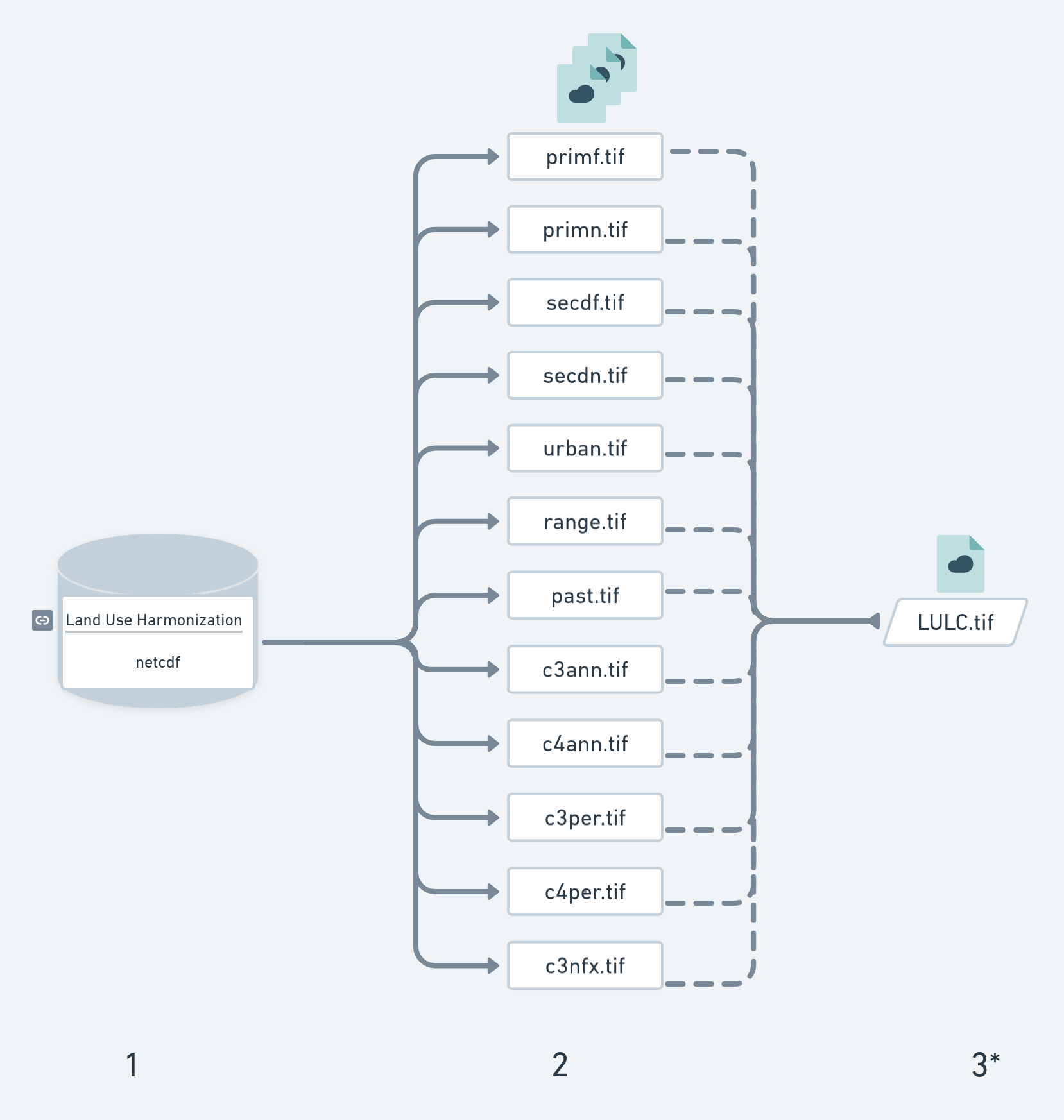

Figure S2. Workflow: 1) Extract every single state per year from 850 to 2100, with two future scenarios (SSP2-4.5 and SSP5-8.5) from 2015 on; 2) Save every state (continuous data) into “State-files” in TIFF format; and  3*) Create new (categorical data) of land-use land-cover combining 12 states into “LULC -iles” in TIFF format.
